# Global early replication disrupts gene expression and chromatin conformation in a single cell cycle

**DOI:** 10.1101/2022.05.11.491470

**Authors:** Miguel M. Santos, Mark C. Johnson, Lukáš Fiedler, Philip Zegerman

## Abstract

The early embryonic divisions of many organisms, including fish, flies and frogs are characterised by a very rapid S-phase caused by high rates of replication initiation. In somatic cells, S-phase is much longer due to both a reduction in the total number of initiation events and the imposition of a temporal order of origin activation. The physiological importance of changes in the rate and timing of replication initiation in S-phase remains unclear. Here we assess the importance of the temporal control of replication initiation using a conditional system in budding yeast to drive the early replication of all origins in a single cell cycle. We show that global early replication disrupts the expression of over a quarter of all genes. By deleting individual origins, we show that delaying replication is sufficient to restore normal gene expression, directly establishing replication timing control in this regulation. Global early replication disrupts nucleosome positioning and transcription factor binding during S-phase, suggesting that the rate of S-phase is important to regulate the chromatin landscape. Together these data provide new insight into the role of a temporal order of origin firing for coordinating replication, gene expression and chromatin establishment as occurs in the early embryo.

## Introduction

Eukaryotic genomes are replicated from multiple start sites or origins. To ensure genome stability, these origins must initiate replication only once during the cell cycle. Strict initiation control first involves the loading of the inactive replicative helicase (Mcm2-7) onto DNA only in late M/G1 phase, in a process called licensing. Initiation at these licensed origins can then only occur during S-phase, due to the activation of S-phase cyclin-dependent kinase (S-CDK) and Dbf4-dependent kinase (DDK). DDK phosphorylates the loaded Mcm2-7 helicase, which drives the recruitment of Sld7/Sld3 (MTBP/Treslin in humans) and the helicase activating protein Cdc45, while CDK phosphorylates Sld3 and another protein Sld2 (RecQL4 in humans), driving interactions with Dpb11 (TopBP1 in humans), DNA polymerases and other factors to form active replisomes and initiate replication [1].

In most eukaryotic cells origins do not all fire at the beginning of S-phase. Although all origins are licensed in G1 phase, only a fraction of these (10% in mammalian cells) will be activated during a normal S-phase [2]. For those origins that fire, their activation occurs as a continuum throughout S-phase, resulting in differences in the timing of genome replication. This temporal order of genome duplication is known as the replication timing (RT) programme. Importantly this RT programme is evolutionarily conserved between related species [3], is consistent within particular cell types [4] and changes during development [5], differentiation [6] and disease [7].

In budding yeast and metazoa, the signal for early or late replication is established in G1 phase during licensing [8, 9]. Several studies suggest that the establishment of RT is dependent on the regulation of the chromatin context of origins [10–13] and early origins differentially bind to the origin recognition complex (ORC) [14, 15] and load more Mcm2-7 [16]. We and others have shown that a critical determinant of origin timing is the ability of licensed origins to compete for a limiting pool of initiation factors [17, 18]. The activatory subunit of DDK, Dbf4, and the two critical CDK targets, Sld3 and Sld2, and their binding partner, Dpb11 are stoichiometrically limiting for initiation in budding yeast and over-expression of these factors, together with the Sld3 partner proteins, Sld7 and Cdc45, is sufficient to allow the early firing of late and dormant origins in yeast [18]. Importantly the vertebrate orthologues of these factors are also rate-limiting for replication initiation during early embryonic divisions [19]. Direct recruitment and inhibition of these limiting initiation factors can also influence replication timing, for example kinetochores and the forkhead box transcription factors Fkh1/Fkh2 can drive the early firing of subsets of origins by directly recruiting Dbf4 [20, 21], while conversely Rif1 counteracts DDK activity to delay RT in late replicating domains [22].

Although differential replication timing of the genome was identified over 50 years ago [23] and RT is dramatically altered during differentiation and development [5, 6], the functional importance of the RT programme is still very poorly understood. Over-expression of limiting replication initiation factors, causing the early firing of many origins in budding yeast, results in cell lethality [18], strongly suggesting that ordered origin firing is important. In many organisms including humans, mouse and *Drosophila* there is a positive correlation between early replication and the probability of gene expression [24], while in budding yeast only the highest expressed genes tend to be earlier replicated while the lowest expressed genes tend to be late replicated [25]. Despite this, whether early replication is a cause or a consequence of gene expression is not clear. Recently, it was shown that perturbations in RT caused by loss of Rif1 in human cells is coupled with alterations in histone modifications and 3D chromatin compartments, but only causes a limited effect on gene expression [26]. In yeast however, early replication of the histone genes is required for their maximal expression in S-phase [27].

Here we take advantage of our system that conditionally over-expresses limiting replication initiation factors in budding yeast [18] to determine the relationship between perturbations in DNA replication timing and gene expression in a single cell cycle. By driving the early replication of the majority of the genome we observe dramatic changes in the expression of over a quarter of yeast genes during S-phase. Early genomic replication causes concomitant changes in chromatin dynamics and transcription factor binding events during S-phase. This study provides new insight into the importance of ordered origin firing, with implications for how replication rate can influence gene expression and the chromatin landscape.

## Results

### Over-expression of limiting replication initiation factors causes global early replication

To perturb replication timing genome-wide in a single cell cycle we took advantage of our conditional system in budding yeast that over-expresses six limiting replication factors Sld2, Sld3, Dpb11, Dbf4, Cdc45 and Sld7 (abbreviated to SSDDCS) upon the addition of galactose [18]. Galactose-induction of the *SSDDCS* strain speeds up S-phase, as measured by flow cytometry, but also leads to the depletion of the dNTP pool, which causes activation of the DNA damage checkpoint [18]. To advance replication timing, without activating the checkpoint we performed all experiments with strains containing a null mutation of the ribonucleotide reductase inhibitor *SML1.* The *sml1*Δ mutation increases the dNTP pool and overcomes the galactose-induced checkpoint activation in the *SSDDCS* strain [18].

To determine the genome-wide impact of over-expression of the limiting replication factors on DNA replication timing in a single cell cycle, we analysed replication progression by DNA sequencing and copy number analysis. For this, the *sml1*Δ and the *sml1Δ SSDDCS* strains were synchronised in G1 using the mating pheromone alpha-factor, galactose was added to these G1 arrested cells for 30 min and then cells were released into a synchronous S-phase in the presence of galactose, whereby samples were collected every 5 minutes to generate libraries for whole genome sequencing (a similar scheme was used for RNA-seq, Figure 2A). To determine the replication dynamics of these strains, the ratio of mapped reads at each timepoint compared to the G1 sample (copy number) was calculated for each genomic 1kb bin, as described [28]. The median replication time (T_rep_) for each 1kb bin was determined by fitting a sigmoidal curve to the copy number profile from each bin and extracting the time (in minutes after G1 release) at which each bin is half-way between one and two copies.

**Figure 1.**
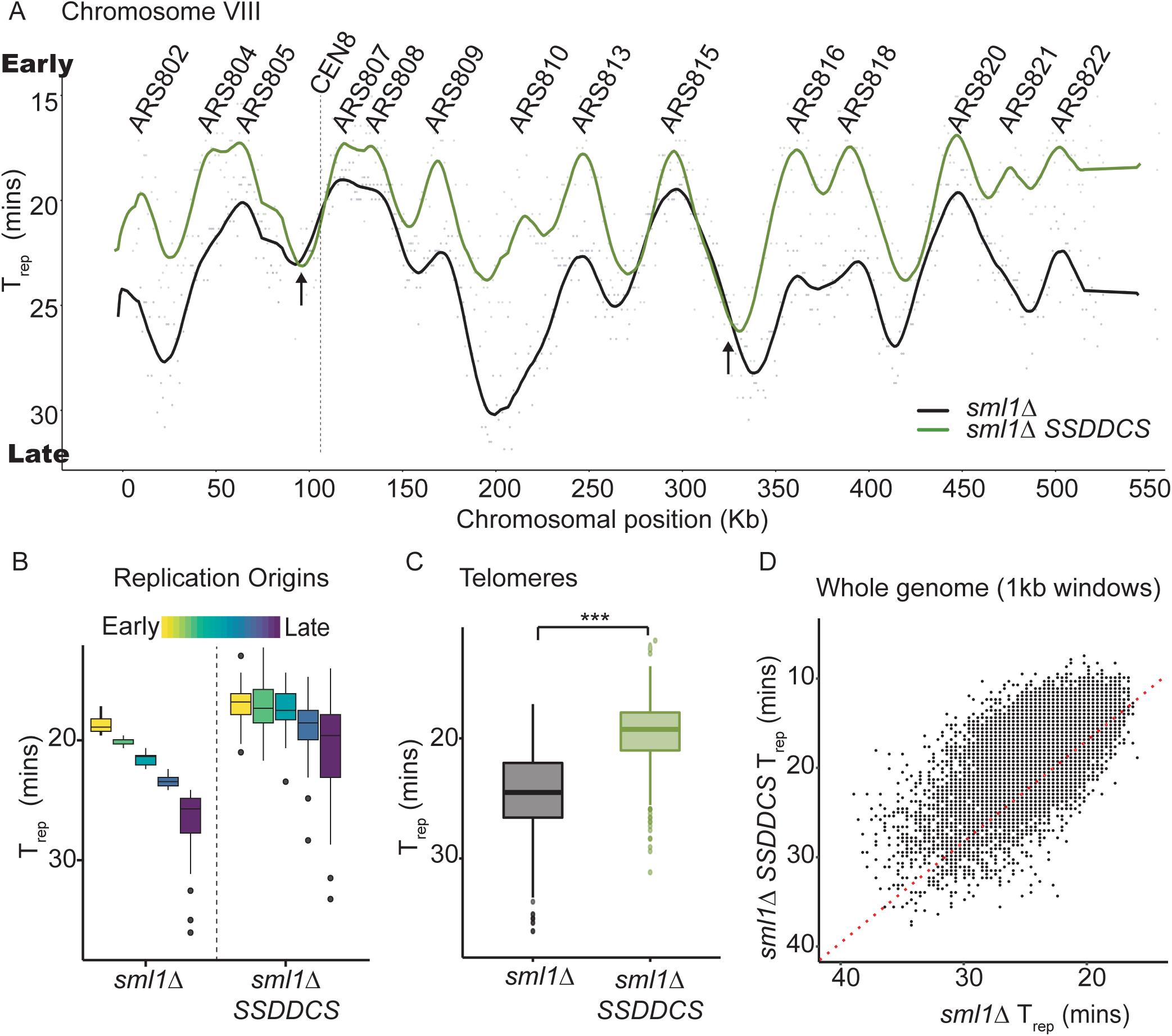
Over-expression of limiting initiation factors causes global early replication. A) T_rep_ values were plotted from the indicated strains along the corresponding chromosome positions to generate genome-wide replication profiles and smoothed using a moving average. Chromosome VIII is shown here as an example. The y-axis is flipped so that early replicating regions are at the top of the plot and late replicating regions at the bottom. The location of annotated origins (ARS) is shown above. Overall, all origins fire earlier in the *SSDDCS* strain. Arrows indicate termination zones that have a delayed T_rep_ in the *SSDDCS* strain. B) Distribution of T_rep_ values for all origins divided into quintiles according to T_rep_ values from the *sml1*Δ strain. C) Distribution of T_rep_ values for all telomeres (genomic bins within 50kb of chromosome ends).*** p-value = < 2.2e-16, Welch Two Sample t-test. D) Scatterplot of the T_rep_ values for the whole genome in 1kb bins. Red dashed line is the line of equal T_rep_ between the *sml1*Δ and *sml1Δ SSDDCS* strains.

**Figure 2.**
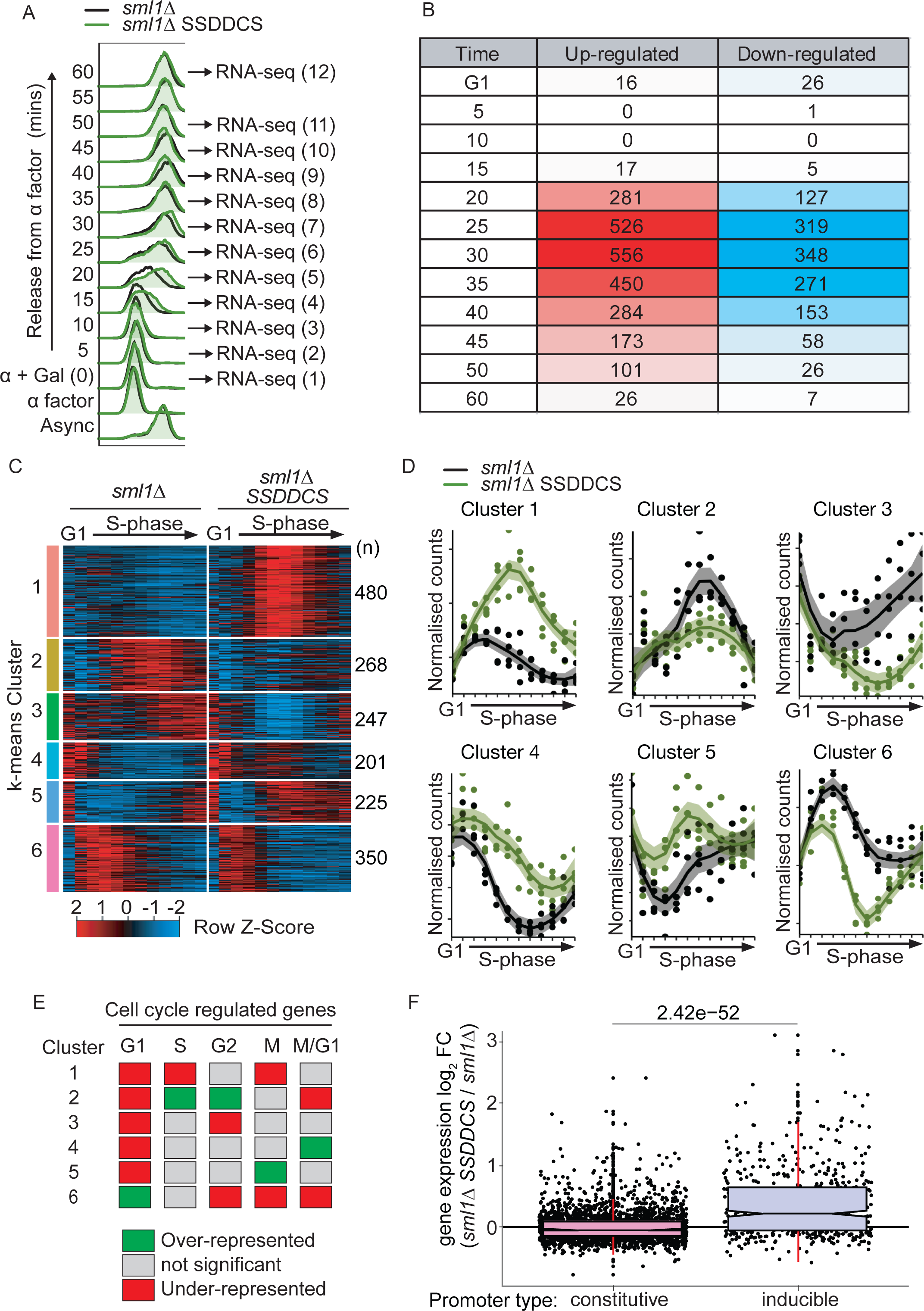
Global early replication perturbs the expression of 27% of genes in one cell cycle. A) Overlay of the flow cytometry between the *sml1*Δ and *sml1Δ SSDDCS* strains arrested in α factor and released into S-phase in galactose. The timepoints (1-12) used to generate time-resolved RNA-seq libraries are indicated. B) Table showing the number of differentially expressed genes at each of the 12 RNA-seq timepoints between the *sml1*Δ and *sml1Δ SSDDCS* strains. Cells are colour-coded based on the number of DE genes on each time-point. C) Heatmap of 1771 genes that are differentially expressed (DE) genes between the two strains in one or more time-points after G1. Each row corresponds to one individual gene and each column to one time-point (G1 to 60 mins from left to right, respectively). Genes were clustered according to their expression profiles using k-means clustering. The expression levels were z-scored and normalised by row. The maximal and minimum expression are coloured in red and blue, respectively. D) Normalised read counts per time-point using DESeq2 size factors for one illustrative gene from each DE cluster from C). E) Table showing k-means clusters from C with the statistically significant (p value <0.05) over- (green) or under- (red) representation of genes designated to be cell cycle regulated according to [33]. p values were calculated using binomial probability tests. F) Box and whisker plot of the log_2_ time-course-averaged expression fold change for *sml1Δ SSDDCS* / *sml1*Δ strains of genes separated into constitutively expressed or inducible according to [34].

Replication profiles were generated by plotting the T_rep_ values along the corresponding chromosome position, followed by smoothing using a moving average (Figure 1A – only Chromosome VIII is shown). From such a profile, the peaks (local minima of T_rep_) represent origins of replication and troughs (local maxima of T_rep_) are termination zones (note that for these analyses the y-axis is inverted so that origins are peaks – Figure 1A). As expected [18, 29], SSDDCS over-expression advances the replication timing of all origins across the whole chromosome (Figure 1A).

To address whether all origins genome-wide were equally affected by over-expression of limiting factors, replication origins were divided into quintiles according to their normal T_rep_ [30] (Figure 1B). This analysis demonstrated that while all origins replicate earlier upon SSDDCS over-expression, late origins have a greater advance in replication timing (Figure 1B). This effect can also be observed at normally late replicating telomeres (Figure 1C), but not at early replicating centromeres (Supplementary Figure 1), which directly recruit limiting replication factors to ensure early replication in budding yeast [21]. Analysis of the whole genome as 1kb bins, reveals that the vast majority of the genome is replicated earlier upon SSDDCS over-expression (Figure 1D, above red line). Despite this, some sites are replicated later after SSDDCS over-expression (Figure 1D, below red line). We have previously shown that high rates of initiation causes topological defects, due to the overwhelming of topoisomerase activities, resulting in delays in termination [31]. Such delayed termination sites are indeed visible on the T_rep_ plots (black arrows, Figure 1A). Apart from these small number of termination zones, the SSDDCS over-expression strain clearly induces the early replication of the majority of the genome in a single cell cycle.

### Global early replication perturbs the transcription of over a quarter of the genome in a single cell cycle

To analyse the impact of advanced replication on gene expression during a single cell cycle, samples were collected for whole transcriptome sequencing (RNA-Seq) from synchronised cells as in Figure 2A. A principal component analysis (PCA) revealed the absence of batch effects between biological replicates (Supplementary Figure 2). To address a role for replication timing in gene expression control, each time-point was compared between the two strains using DESeq2 [32]. Note that due to the over-expression of the six factors in the *SSDDCS* strain, these six genes were excluded from all downstream analyses. Genes were considered up or down-regulated in the *SSDDCS* strain if the log_2_ normalised fold-change (*SSDDCS sml1Δ* / *sml1*Δ) was above or below 0 respectively and if the adjusted p-value or false discovery rate (FDR) was below 0.01 (DESeq2 Wald test). Using this approach, a large number of genes were differentially expressed (DE) in the *SSDDCS* strain in a single cell cycle (Figure 2B). Importantly, the majority of these expression changes occur after 20 minutes, suggesting that S-phase is required for these differences (Figure 2B). Notably the number of DE genes was greatly reduced by the end of the time course, suggesting that any changes in expression are resolved before the next cell cycle (Figure 2B).

Taking advantage of the temporal resolution of this dataset, we analysed the expression pattern of each gene during the cell cycle, using the DESeq2 likelihood ratio test (LRT) [32]. A significant FDR from this test identifies genes that show a difference between the two strains at one or more time-points after G1 phase. Using this approach, a total of 1771 genes were identified as differentially expressed, representing ∼27% of the genome. From this list, k-means clustering was used to group genes with similar expression profiles (Figure 2C/D). This clustering predominantly segregated genes by their normal periodic expression [33] (Figure 2E), for example Cluster 2 genes are significantly over-represented in S/G2 expressed genes. As the k-means clusters indicated that the genes that are differentially expressed in the *SSDDCS* strain are dynamically regulated even in a wild type strain (Figure 2D), we wondered whether constitutively expressed genes, which do not change their expression under different conditions [34], might be less affected than inducible genes by advanced replication timing. Figure 2F shows that constitutively expressed genes show much less differential expression than inducible genes after early replication, consistent with the fact that these constitutive genes are refractory to environmental stimuli [34].

We wondered to what extent replication timing changes might be important for the gene expression changes that we observed (Figure 2B-D). As SSDDCS over-expression causes the early replication of the majority of the genome (Figure 1D), it was not surprising that all DE gene clusters, as well as non-DE genes are on average earlier replicated in the *sml1Δ SSDDCS* strain compared the *sml1*Δ strain (Supplementary Figure 3A). Analysis of the proximity of DE genes to origins revealed that clusters 1,4 and 5 are on average more origin proximal than non-DE genes, and vice versa for the remaining clusters (Supplementary Figure 3B). Clusters 1 and 4 show enrichment for genes that are proximal to the normally late-replicating telomeres, although the majority of DE genes are not telomeric (Supplementary Figure 3C). Interestingly, given that centromeres do not show replication timing changes in the *SSDDCS* strain (Supplementary Figure 1), they are also not enriched for DE genes from any of the k-means clusters (Supplementary Figure 3D). This data suggests that advanced replication timing is likely to be necessary for the gene expression changes that we observe in the *SSDDCS* strain, but advanced timing is not sufficient for these changes as non-DE genes are also earlier replicating in the *SSDDCS* strain (Supplementary Figure 3A).

The expression of some genes is directly affected by an increase in copy number during replication, such as the histone genes [27]. It has also been suggested that to maintain expression homeostasis during S-phase, the earliest S-phase genes are buffered against copy number changes through the histone acetyltransferase Rtt109 [35]. As early replication of the genome in the *SSDDCS* strain might affect gene expression by increasing the copy number of a gene in early S-phase, we wondered to what extent this might influence the differential expression we observe during advanced replication (Figure 2C). Importantly, we observed no correlation between the genes that are DE in the *SSDDCS* strain and the *rtt109Δ* sensitive or insensitive gene sets [35] (data not shown). In addition, the k-means clusters that are enriched for genes that are normally expressed in S-phase and G1/S phase (clusters 2 and 6 respectively, Figure 2E) show reduced expression after early replication (Figure 2C/D), which is the opposite of what we would expect if copy number was affecting their expression. Finally, the differences in expression between *sml1*Δ and *sml1Δ SSDDCS* were significantly greater than 2-fold for many genes (in particular for Cluster 1, e.g see Figure 3A), again suggesting that the expression changes of these genes are not simply due to copy number changes. Together these data show that genome-wide early replication perturbs the normal expression of a large proportion of the genome in a single cell cycle.

**Figure 3.**
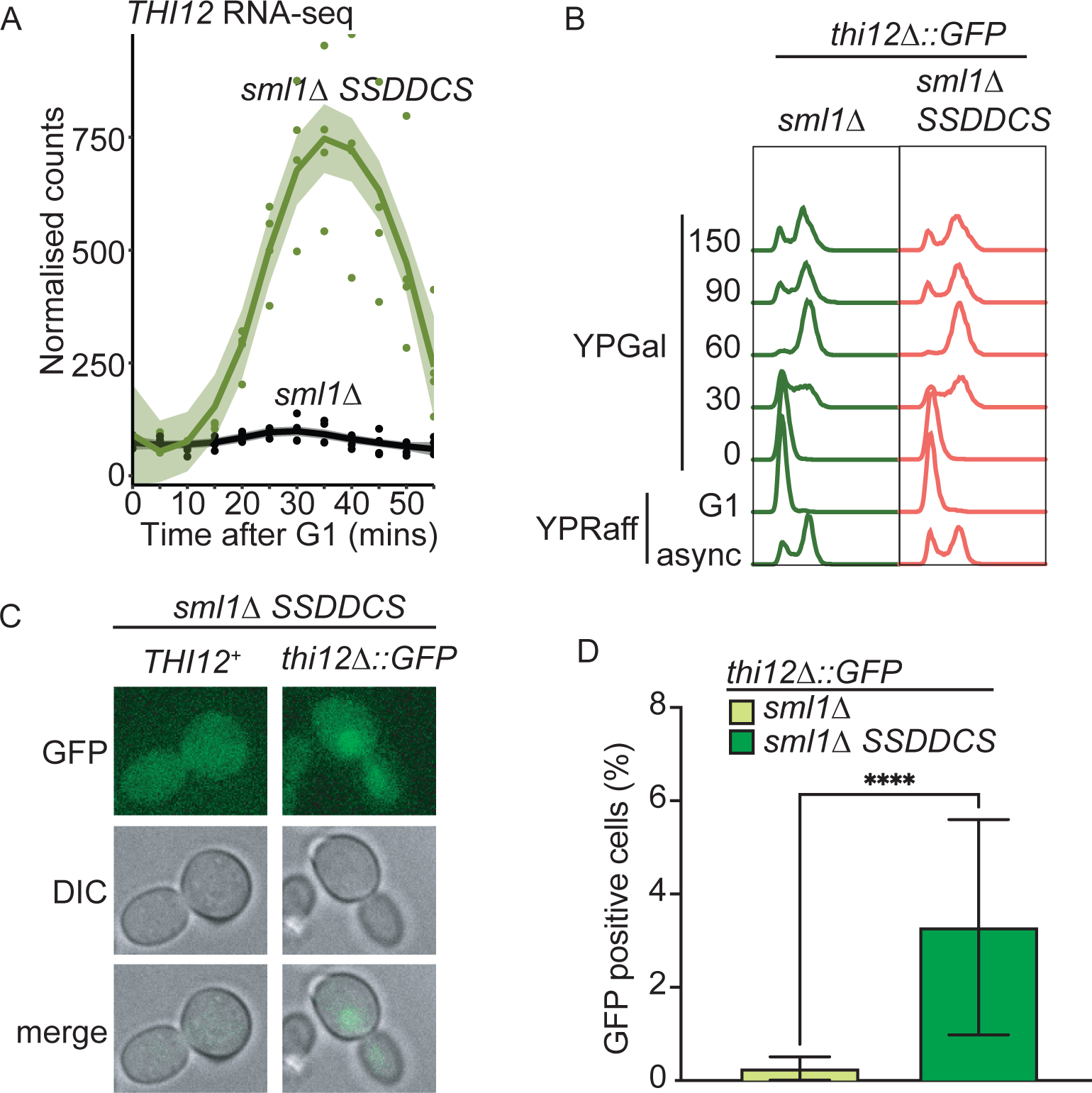
S**ingle cell analysis of early replication-induced gene expression** A) RNA-seq data for *THI12,* a cluster 1 DE gene, from the data in Figure 2C. B) Flow cytometry of the timecourse for analysis of GFP expression from the *THI12* locus. C) Representative images from the analysis of GFP expression at the 150 mins timepoint. D) Quantitation of GFP expression from the 150 mins timepoint as in C for 3 independent experiments. n=755 and 1156 for *sml1*Δand *sml1Δ SSDDCS* respectively. Error bars are SD, p-value <0.001 from an unpaired T-test.

### Single cell analysis of early replication induced transcription

From the RNA-seq analysis we can conclude that altering replication timing affects the expression of many genes across a population of synchronised cells. We wondered whether the effect of replication timing on gene transcription was of sufficient magnitude to observe these expression changes in individual cells in a single cycle. For this, we replaced the *THI12* gene with *GFP* as a fluorescent proxy for *THI12* expression. *THI12* is a cluster 1 gene that is highly expressed in the *SSDDCS* strain (Figure 3A). We replaced *THI12* with a version of GFP that has a PEST degron sequence from Cln2 to ensure that the protein is short-lived and an NLS to concentrate the protein in the nucleus and enhance the microscopy analysis [36]. We performed the same experiment as with the RNA-seq (Figure 3B), except that we took later timepoints, due to the delay between GFP expression, translation and folding [36]. Using this system, we observed that GFP was not expressed in the *sml1*Δ strain in galactose as expected, as *THI12* is not expressed in these conditions (Figure 3A/D). However, in the *SSDDCS* strain GFP was significantly expressed in approximately 3% of cells in a single cell cycle (Figure 3C/D). Although only a small percentage of cells appeared positive, we do not know what threshold of GFP expression is necessary for a positive signal in this assay. Despite this, it is clear that early replication indeed affects gene transcription in individual cells in a single cell cycle.

### Delaying replication rescues aberrant transcriptional activation

If early replication is required for the aberrant gene expression that we observe (Figure 2), then we predicted that if we could delay the replication of individual genes in the *SSDDCS* strain, we might be able to restore their normal expression. For this we identified two cluster 1 genes, *IME2* and *NDT80,* that are highly expressed in S-phase in the *SSDDCS* strain, similar to *THI12* (Figure 3A) and are also proximal to active origins (Figure 4A/B). Taking advantage of the fact that replication origins are defined genetic elements in budding yeast (autonomously replicating sequences - ARS), we deleted the two neighbouring ARS elements from the *IME2* locus (ARS1008/1009, Figure 4A) and from the *NDT80* locus (ARS816/818, Figure 4B). Importantly these ARS deletions specifically delayed the replication of the target locus (Figure 4A/B), but the replication timing of all other origins remained similar between the strains (Supplementary Figure 4). Analysis of the expression of the *IME2* and the *NDT80* genes by qPCR showed that, as expected these genes are not expressed in the *sml1*Δ strain but are expressed in S-phase in the *sml1Δ SSDDCS* strain (Figure 4C/D). Significantly, the deletion of the neighbouring origins, which delays replication specifically of the *IME2* or the *NDT80* loci, completely abrogated the aberrant expression of these two genes in the *SSDDCS* strain (Figure 4C/D). This experiment strongly suggests that it is the early replication of these loci that is critical for their aberrant S-phase expression.

**Figure 4.**
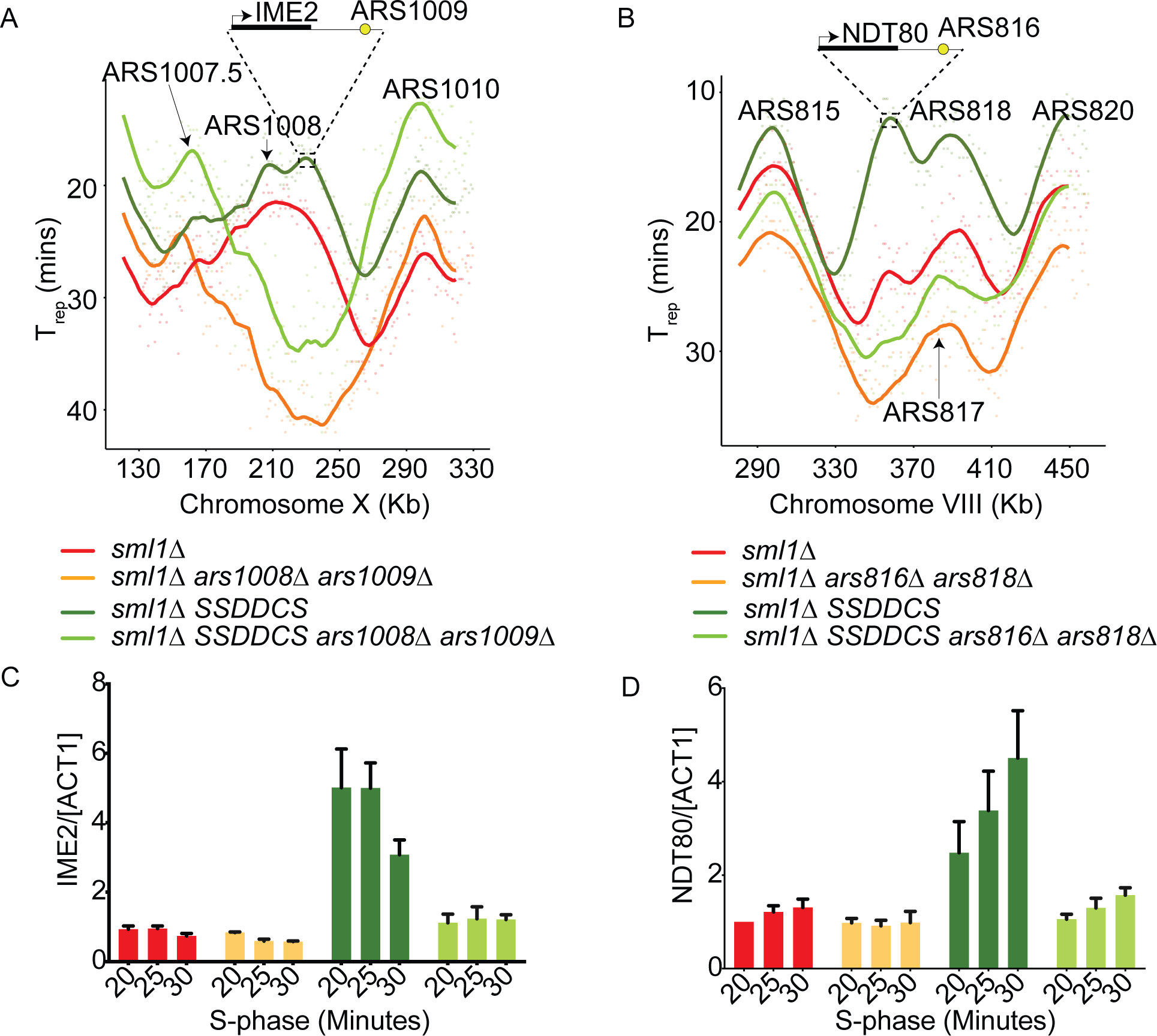
Delaying replication rescues aberrant transcriptional activation. A) T_rep_ profile from the indicated strains as in Figure 1A, from a segment of Chr X. The position of the *IME2* locus is indicated on the *sml1Δ SSDDCS* profile. B) As A, except for the *NDT80* locus on Chr VIII. C) qPCR data of *IME2* expression from the indicated timepoints for the strains as in A. Error bars are SD, n=3. Gene expression was normalized to actin. D) As C, except for *NDT80* expression.

### Global early replication perturbs chromatin conformation in a single cell cycle

To address how galactose induction of the *SSDDCS* strain might lead to dramatic changes in transcription in a single cell cycle, we set out to analyse whether early replication of the genome affects chromatin organisation. For this, we analysed nucleosome positioning and occupancy across the genome by sequencing after micrococcal nuclease digestion (MNase-seq). Most genes contain a well-positioned nucleosome (+1 nucleosome), just downstream of the transcriptional start site (TSS), which is dynamically repositioned during transcription [37, 38]. From our MNase-seq data, we observed that the position of the +1 nucleosome remains largely static relative to the TSS in the *sml1*Δstrain, regardless of whether the genes are expressed or not during the time course (Figure 5A). However, in the *SSDDCS* strain that induces global early replication, all DE and even non-DE genes displayed greater mobility of the +1 nucleosome towards the gene body (Figure 5A). Importantly this mobility of the +1 nucleosome was transient and was restored to the normal position by the end of the time course (Figure 5A).

**Figure 5.**
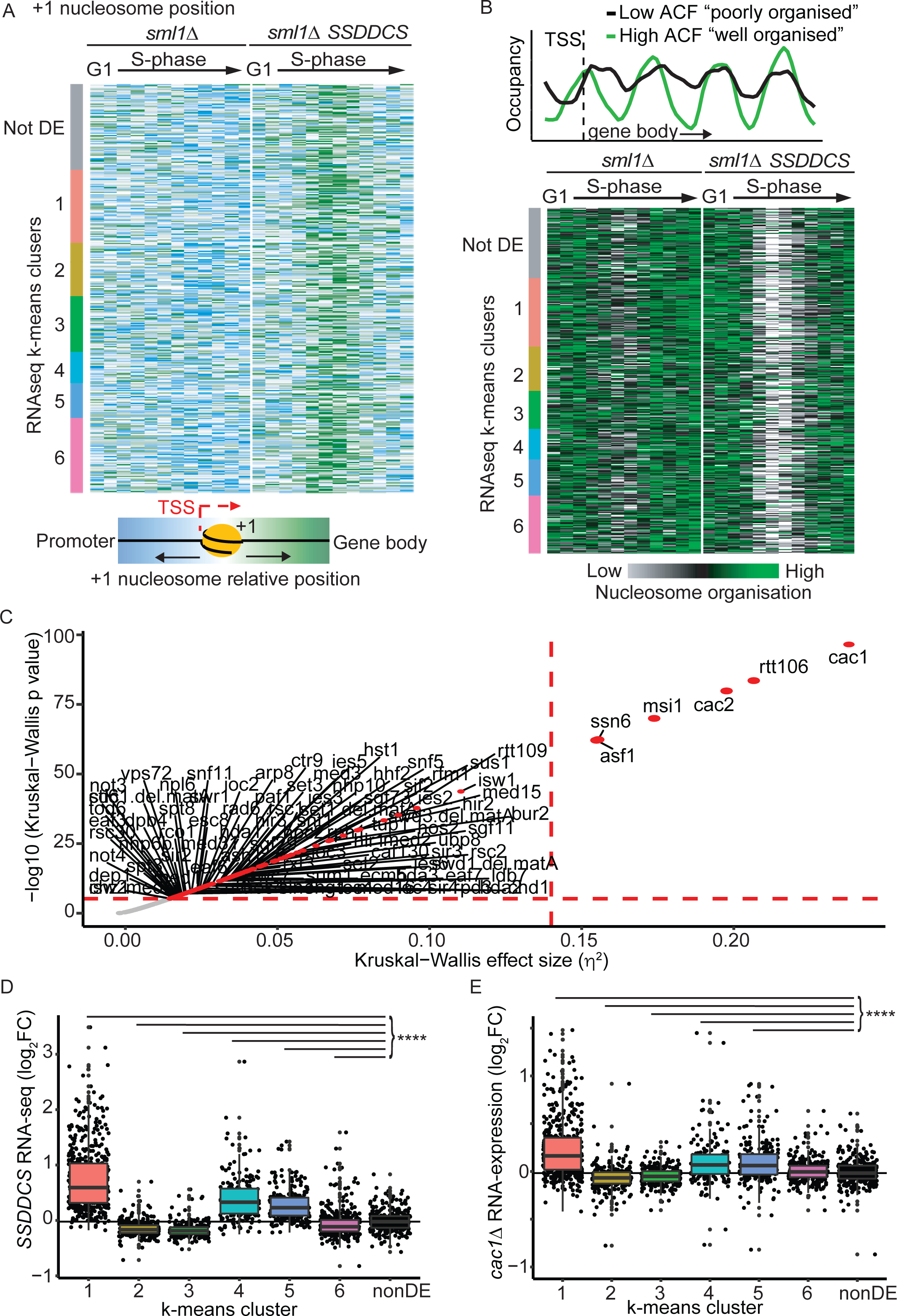
Global early replication perturbs chromatin conformation in a single cell cycle. A) Position of the +1 nucleosome relative to the transcriptional start site (TSS) represented as a heatmap for the DE genes separated into their k-means cluster (Figure 2C) plus 250 random non-DE genes with an annotated +1 nucleosome. Blue represents the most upstream position of the +1 nucleosome, while green represents the most downstream position (into the gene body). Data is normalised by row as done for previous heatmaps. Each row corresponds to a single gene and each column to a single time-point (G1 to 60 from left to right, respectively). B) Top – The nucleosome occupancy and organisation of the first 4 nucleosomes after the TSS was analysed using an autocorrelation function (ACF) analysis: nucleosomes in green are well phased (i.e. well defined peak and valleys) and well positioned (similar inter-nucleosome distance), corresponding to high ACF values. On the other hand, nucleosomes in black are poorly phased and disorganised, resulting in low ACF. Bottom - Heatmap of ACF values for the DE genes and 350 random non-DE genes longer than 700bp. Each row corresponds to a single gene and each column to a single time-point (G1 to 60 from left to right, respectively). The two strains are separated by a white vertical line. White and green represents the time-points at which the chromatin is mostly disorganised and organised, respectively. C) Plot of statistical significance versus effect size for Kruskal-Wallis tests applied to chromatin mutant gene expression data from [40]. Red dashed horizontal and vertical lines represent a Bonferroni-corrected p value threshold of 0.001 and a high-effect threshold (η^2^ = 0.14), respectively. D) Box and whisker plot of the averaged gene expression change in the SSDDCS strains for the k-means clusters from the RNA-seq data in Figure 2C. p values are from Holm- corrected Wilcoxon rank sum tests relative to 300 random non-DE genes (black). **** p < 0.0001. E) Box and whisker plot of the RNA expression changes in the *cac1Δ* strain (data from [40]). p values are from Holm-corrected Wilcoxon rank sum tests relative to 300 random non-DE genes (black). **** p < 0.0001.

To analyse nucleosome organisation in the gene body, the pattern of the first 4 nucleosomes was assessed using the autocorrelation function (ACF), as recently described [39]. Genes with higher ACF values have well-phased and organised nucleosomes, while lower values represent poorly organised chromatin (Figure 5B, top). ACF values were calculated for all genes in each time-point and plotted as a heatmap (Figure 5B, bottom). This analysis was restricted to genes larger than 700bp (as 4 nucleosomes are needed for ACF calculation), which corresponds to approximately 70% of all genes. In the *sml1*Δ control strain there is a transient decrease in nucleosome organisation during S-phase, most likely caused by the passage of replication forks, but once replication is completed the nucleosome organisation is re-established (Figure 5B). Strikingly, a greater decrease in nucleosome organisation was observed genome-wide in the *SSDDCS* strain (Figure 5B), which is in agreement with the +1 nucleosome data (Figure 5A). This decreased nucleosome organisation in the *SSDDCS* strain was not specific to the DE genes and was restored back to normal levels of organisation by the end of the time course (Figure 5B). Together this data shows that genome-wide early replication perturbs the nucleosome positioning and organisation of many genes.

Figure 5A and 5B demonstrate that driving early replication genome-wide perturbs chromatin organisation during S-phase. One mechanism that might cause this defect could be that the high rate of replication early in S-phase overwhelms the histone supply and assembly machinery, leading to a transient period when new nucleosome deposition/positioning cannot keep pace with replication. If such transient chromatin assembly defects are responsible for the global gene expression changes that we observe in the *SSDDCS* strain, then it would follow that strains that are defective in chromatin assembly should mimic the gene expression changes in the *SSDDCS* strain. To investigate the relationship between chromatin defects and expression of the *SSDDCS*-dependent genes, we compared our data with the gene expression changes in a compendium of 165 chromatin machinery deletion mutants, which includes nucleosome remodellers, histone chaperones, histone modifiers and transcription co-regulators [40]. As the gene expression data from these chromatin machinery deletion datasets represent only a single asynchronous time point, we converted our gene expression data (Figure 2C) to an average gene expression value for the entire time course (Figure 5D). Importantly, even after averaging the gene expression time course data, each k-means cluster remained highly significantly different to the non-differentially expressed control group (Figure 5D). This data was then used to quantify the similarity between the *SSDDCS* data and the 165 chromatin machinery mutants using Kruskal-Wallis tests. A plot of statistical significance against effect size clearly identified six mutants with highly significantly similar expression profiles to our *SSDDCS* data (Figure 5C): the CAF-1 histone chaperone complex (Cac1, Cac2, Msi1), the Rtt106 and Asf1 histone chaperones, and the Ssn6 transcriptional co-repressor. Direct comparison of the *cac1*Δ dataset with our *SSDDCS* data mimicked the significant upregulation of the clusters 1/4/5 and the downregulation of clusters 2/3 (Figure 5D versus 5E). The magnitude of the fold change in the *cac1*Δ data was smaller than for our *SSDDCS* data (Figure 5D versus 5E), but this might reflect methodological differences from comparing our RNA-sequencing data with the *cac1*Δ microarray data from asynchronous mid-log yeast populations [40]. Together, the dramatic defect in chromatin organisation in mid-S-phase (Figure 5A/B) combined with the correlation of our gene expression data with chromatin assembly mutants (Figure 5C-E) strongly suggest that early replication of the entire genome causes a transient chromatin assembly defect, which may explain to some extent the changes in gene expression that we observe.

### High rates of initiation disrupt the TF landscape

Although there is a correlation between chromatin disruption and changes in gene expression (Figure 5), it is not clear why changes in chromatin should affect the expression of certain genes. Newly synthesised DNA must reassemble nucleosomes and transcription factors (TFs), which compete for binding to nascent DNA [41]. We therefore wondered whether chromatin disruption caused by global early replication, might also perturb the TF landscape. Analysis of nucleosome positioning at the *NDT80* locus (Figure 6A), which is a cluster 1 gene that is highly expressed in S-phase in the *SSDDCS* strain (Figure 4), shows not only that the +1 nucleosome moves into the gene body during S-phase, as expected (Figure 5A), but that the nucleosome positions around the binding sites for the key *NDT80* regulatory TF Ume6 also change during S-phase, specifically in the *SSDDCS* strain (Figure 6A). To address whether nucleosome positioning was altered at other Ume6 binding sites (URS1 sites), these sites were extracted from the JASPAR database [42] and the locations of all URS1 sequences in the budding yeast genome were identified using Find Individual Motif Occurrences (FIMO) [43]. Of the 2875 URS1 sequences identified, 89 were located 1kb upstream of the TSS of genes from cluster 1. Analysis of the average nucleosome profile centred around these 89 locations, showed that nucleosome positioning and occupancy was identical between the strains in G1 phase (Figure 6B, top). In S-phase however, the *SSDDCS* strain exhibited reduced occupancy of nucleosomes at the Ume6 binding sites and movement of the nucleosomes away from these sites (Figure 6B). This suggests that when chromatin is perturbed by global early replication the interplay between TF binding and nucleosome positioning also changes.

**Figure 6.**
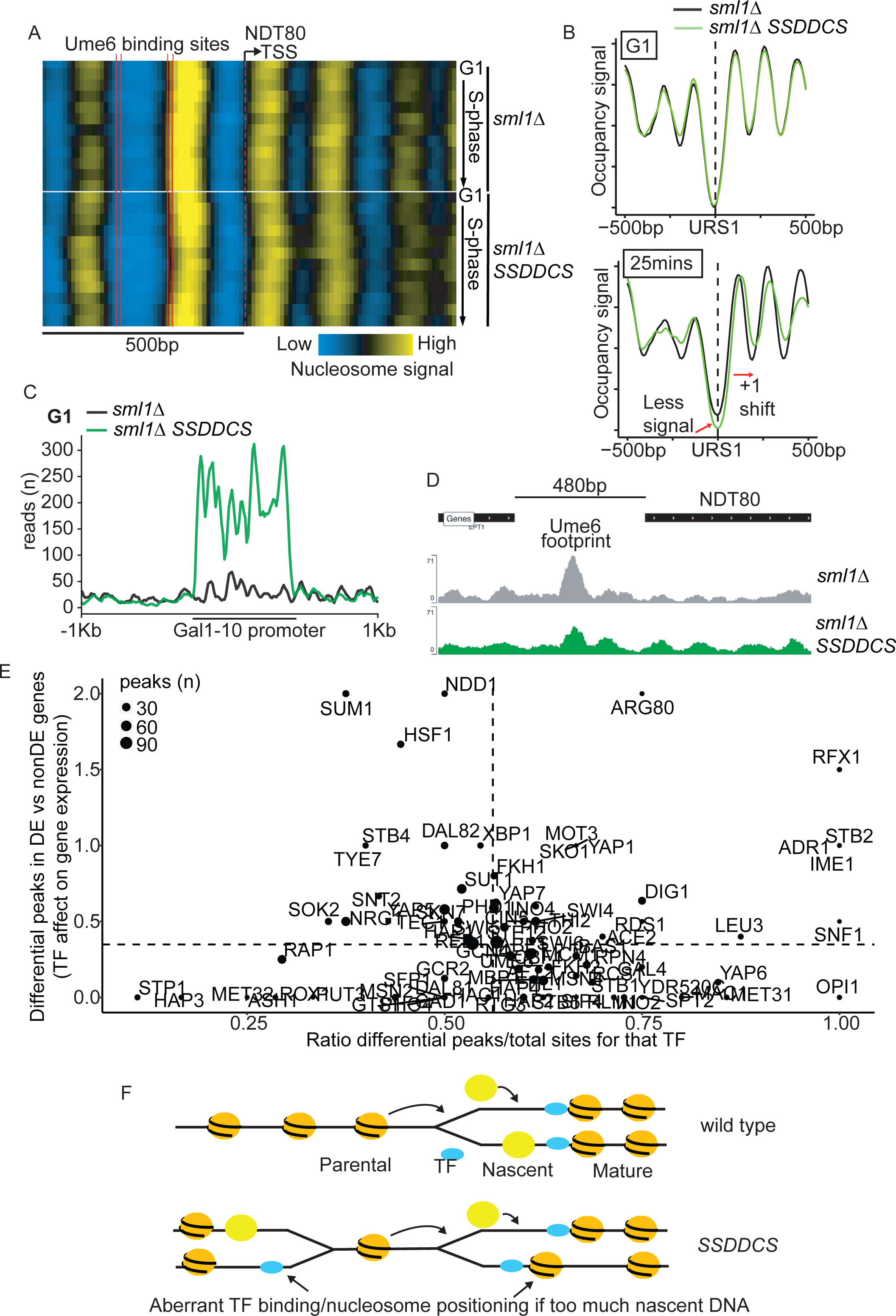
High rates of initiation disrupt the transcription factor landscape. A) Nucleosome positioning heatmaps of the *NDT80* locus, which has two Ume6 binding sites in the promoter region (vertical solid red lines). The vertical dashed red line marks the TSS and each row represents one time-point, from G1 to 60 from top to bottom. Yellow corresponds to a high density of MNase-seq reads corresponding to nucleosome peaks, while blue corresponds to nucleosome depleted regions. B) Average nucleosome profiles for the 89 URS1 locations identified in promoters of cluster 1 genes in G1 and at 25 minutes. The profiles are centred on the URS1 sequence (the Ume6 binding site - dotted line). C) Sub-nucleosomal MNase-seq read coverage around the GAL1-10 promoter in G1. D) Sub-nucleosomal MNase-seq reads of the *NDT80* locus 5 minutes after release from G1 phase. Y-axis shows the sub-nucleosomal read coverage in this genomic region. Top track shows the location of genes. Peak annotated to the Ume6 binding site is highlighted. E) Each sub-nucleosomal MNase-seq peak was assigned to a known TF binding site from [44]. For each TF, the ratio of peaks that change in the *SSDDCS* strain over total number of peaks for that TF (x-axis “TF binding changes”) was plotted against the ratio of the differential TF peaks that are in the promoters of genes that are DE or non-DE in the *SSDDCS* strain (y-axis “TF effect on gene expression”). The size of the points is proportional to the total number of sub-nucleosomal peaks identified for each TF. The dashed lines mark the ratios when considering all sub-nucleosomal peaks. F) In wild type cells, nascent DNA is loaded with old and new nucleosomes (yellow) in competition with TFs (blue) as the replication fork progresses. Post-replicative chromatin matures via a variety of mechanisms. In the *SSDDCS* strain, which drives global early replication, lots of concurrent replication forks increase the rate of nascent DNA production, altering the balance of competition between nucleosomes and TFs, leading to aberrant chromatin organisation and TF positioning. This increased S-phase rate provides a window of opportunity to change the pattern of gene expression in a single cell cycle.

To analyse TF binding dynamics in an unbiased way, we used an optimised version of MNase-Seq that allows the analysis of DNA binding proteins such as TFs that are below the size of single nucleosomes [39]. Using this approach and analysing only fragments smaller than 100bp, we identified 7493 sub-nucleosomal peaks. To pinpoint differential TF binding events that could explain the observed differences in gene expression, only peaks that were located 1kb upstream of a TSS and that had a 2-fold difference in the *SSDDCS* strain in at least 1 time-point outside G1 phase were selected (2950 peaks). These 2950 sites represent 61% of all the peaks that we identified in gene promoters suggesting that early replication affects a majority of TF binding events during S-phase.

In order to verify that the sub-nucleosomal peaks we identified correspond to genuine TF binding events, peaks were mapped to annotated TF binding sites [44]. Only peaks within 200bp of an annotated site were considered for downstream analyses (1538 peaks). As the *SSDDCS* strain contains extra copies of the *GAL1-10* promoter (which are used to overexpress the six factors), we analysed TF binding to this promoter to confirm that the subnucleosomal MNase-seq isolated genuine TF binding locations. As expected, the read coverage at the *GAL1-10* promoter was significantly higher in the *SSDDCS* strain compared to the *sml1*Δ strain (Figure 6C), suggesting that TF binding events can be identified using this method. In line with the nucleosome mapping data (Figure 6A/B), we also observed changes in sub-nucleosomal peak abundance in the *SSDDCS* strain in the promoter region of *NDT80*, which overlap with the Ume6 binding sites (Figure 6D). As Ume6 represses *NDT80* expression in mitotic cycles [45], this reduced Ume6 footprint (Figure 6D) fits with the increased *NDT80* expression in the *SSDDCS* strain (Figure 4). To analyse all sub-nucleosomal peaks and their relationship to changes in gene expression in the *SSDDCS* strain we plotted the fraction of differential peaks for a particular TF (as a ratio of that TF’s total number of genomic sites, x-axis Figure 6E) versus the number of those differential peaks that occur in the promoters of DE genes (y-axis, Figure 6E). These two ratios allow the simultaneous comparison of the effect of advancing replication timing on TF binding (x-axis) and gene expression for each particular TF (y-axis). From this analysis we observe that the binding of many TFs to gene promoters is affected in the *SSDDCS* strain that advances replication timing (Figure 6E). Despite this, altered TF binding is not sufficient for gene expression changes, as differential TF binding also occurs in the promoters of genes that are not differentially expressed after advanced replication timing (Figure 6E, below dotted line).

## Discussion

Here we have used a conditional system to overexpress six limiting initiation factors in budding yeast and advance replication timing genome-wide in a single cell cycle. This global advance in replication timing was accompanied by a change in the transcription of approximately 27% of all genes (Figure 2), as well as by dramatic changes in chromatin organisation (Figure 5) and transcription factor binding (Figure 6) during S-phase. A potential explanation for these data is that the high rate of replication initiation induced in the *SSDDCS* strain causes a defect in nucleosome positioning/occupancy because the chromatin assembly machinery cannot keep pace with the rapid accumulation of nascent DNA (Figure 6F). Importantly, mutation of the chromatin assembly factor Caf1 (*cac1Δ*) not only mimics the gene expression changes caused by global early replication (Figure 5C-E), but also mimics the reduced occupancy and altered positioning of nucleosomes on nascent chromatin (Figure 5A/B) [46]. From this it is likely that a significant fraction of the transcriptional changes that we observe are therefore a consequence of the chromatin perturbations that arise from defects in nucleosome deposition on nascent DNA [46]. This does not exclude that some genes may have altered expression directly due to replication timing changes via other mechanisms, but such genes may be masked by the global defects in chromatin assembly.

Transcription factors and nucleosomes compete for binding to nascent DNA [41]. Since we show that global early replication causes dramatic changes in chromatin organisation, it is not surprising that we also identify significant changes in TF dynamics (Figure 6). From our experiments we cannot determine whether TF dynamics are affected by chromatin perturbation or vice versa. It is clear however that maintaining the balance between replication rate and re-assembly of the chromatin and TF landscape on nascent DNA is likely to be an important aspect of why origin firing is distributed throughout S-phase in wild type cells (Figure 6F) and may be an explanation for why the *SSDDCS* strain loses viability in galactose [18].

A striking feature of our experiments is that the massive perturbations of chromatin and gene expression that we induce with global early replication, return to normal after S-phase (Figure 2 and Figure 5). Although deposition of new histones is replication fork-dependent, the maturation of chromatin and the re-establishment of nucleosome/TF positioning is dependent on post-replicative factors such as chromatin remodelling [47]. As we performed our experiments in a single cell cycle, it seems that despite the dramatic replication-dependent perturbations in chromatin and TF binding that we induce in the *SSDDCS* strain, the post-replicative maturation pathways can restore the normal regulatory landscape before mitosis. Significantly, post-replicative chromatin maturation pathways also restore the normal nucleosomal landscape in *cac1Δ* mutants, which mimic the chromatin and gene expression changes we observe in the *SSDDCS* strain (Figure 5) [46]. Although the rate of chromatin maturation differs between loci [39], we did not detect any correlation between the genes that are differentially expressed during early replication and sites of rapid or slow maturation kinetics (data not shown). In addition, delaying replication at individual loci in the *SSDDCS* strain by deleting local origins (Figure 4) was sufficient to restore normal expression of these loci suggesting that it is the transient perturbation of chromatin positioning/TF binding in mid-S-phase caused by global early replication (Figure 5) that creates the permissive window for changes in gene expression. This fits well with previous work showing that the complete absence of replication causes very little change in gene expression during S-phase [48], as we would predict that the lack of replication also removes the window of opportunity for chromatin/TF binding changes on nascent DNA (Figure 6F).

The potential for increased rates of replication to create a window of opportunity for changes in gene expression described in this study has implications for understanding the phenotypic changes in cells in which the rate of replication is regulated or mis-regulated. For example, during erythroid differentiation the shortening of S-phase favours changes in cell fate [49]. In addition, during senescence extra origins become activated [50] and senescent cells experience large-scale changes in chromatin organisation and gene expression [51, 52]. As oncogene-activation can also drive senescence, increased origin firing rates and chromatin/gene expression changes [24, 53], this study may provide a new mechanistic link between changes in the rate of replication and the chromatin/gene expression perturbations in cancers.

The embryonic divisions of many organisms, such as fish, flies and frogs, are associated with very high rates of replication initiation, but these early cell divisions lack zygotic transcription and the chromatin landscape remains immature during this period [54].

Maintaining high rates of replication delays the normal timing of genome activation [19], but the direct role of replication in the establishment of the chromatin landscape and the transcriptionally competent state of the zygote is poorly understood [55]. This study provides new links between replication control and gene expression changes, which may be relevant to understand how key factors such as Rif1 coordinate the events of early embryonic development, through regulating both chromatin organisation and replication timing [22, 56].

## Acknowledgements

We thank members of the Zegerman lab for comments and technical support. We also thank Charles Bradshaw (Gurdon Institute, Cambridge, UK) for bioinformatic support, Kay Harnish (Gurdon Institute) for sequencing and the Gurdon Institute imaging facility. We are grateful to Naama Barkai (Weizmann Institute of Science, Israel), Yoav Voichek (Gregor Mendel Institute of Molecular Plant Biology, Austria) and Craig Peterson (University of Massachusetts Medical School, USA) for sharing data and we thank Monica Gutierrez and David MacAlpine (Duke University, USA) for sharing protocols and advice for MNase-seq analysis. Work in the PZ lab was supported by AICR 10-0908, Wellcome Trust 107056/Z/15/Z, Cancer Research UK C15873/A12700 and Gurdon Institute funding (Cancer Research UK C6946/A14492, Wellcome Trust 092096). MS was funded by the BBSRC BB/M011194/1 and a Cambridge Trust scholarship.

## Methods

### Yeast strains

All strains are *W303a ade2-1 ura3-1 his3-11, 15 trp1-1 leu2-3, 112 can1-100 rad5-535*

| Strain | Relevant genotype |
| --- | --- |
| PZ356 | *sml1Δ::URA3* |
| PZ523 | *leu2::Sld7-P<sub>GAL1-10</sub>-Cdc45::LEU2 his3::SLD3-A- P<sub>GAL1-10</sub>- Dbf4-A::HIS3 trp1::Sld2-P<sub>GAL1-10</sub>-Dpb11::TRP1 sml1Δ::HphMX* |
| PZ1407 | As strain PZ523, but *ARS1008Δ ARS1009Δ* |
| PZ1435 | As strain PZ356, but *ARS1008Δ ARS1009Δ* |
| PZ3004 | As strain PZ356, but *ARS816Δ ARS818Δ::KanMX* |
| PZ3005 | As strain PZ523, but *ARS816Δ ARS818Δ::KanMX* |
| PZ4505 | *sml1Δ::HphMX thi12Δ::GFP-PEST-NLS ADE2<sup>+</sup>* |
| PZ4504 | *leu2::Sld7-P<sub>GAL1-10</sub>-Cdc45::LEU2 his3::SLD3-A- P<sub>GAL1-10</sub>- Dbf4-A::HIS3 trp1::Sld2-P<sub>GAL1-10</sub>-Dpb11::TRP1 sml1Δ::HphMX thi12Δ::GFP-PEST-NLS ADE2<sup>+</sup>* |

### Block and release time-course

The *sml1*Δ and *sml1*Δ *SSDDCS S.cerevisiae* strains were grown overnight in YP-raffinose at room temperature. After ensuring that cultures were growing exponentially, 100ml was transferred to 30°C shaking water bath for one cell cycle. At 1x10^7^ cells/ml, 90μl of stock solution of alpha factor was added to 100ml of culture (1:900 dilution) and after 90 minutes 45μl of stock solution of alpha factor (5mg/ml) was added. To confirm the G1 arrest, cells were analysed under microscope using a haemocytometer and the arrest was considered successful if more than 95% of cells had the G1 characteristic shape (“shmoo”) or were unbudded. Upon arrest, 10ml of 20% galactose was added to the cultures to induce the over-expression of the six factors. 30 minutes post galactose addition, G1 samples were collected. Then cultures were washed twice with fresh YP-galactose to release cells from G1 arrest and resuspended in 100ml of YP-galactose. Cultures were maintained at the 30°C shaking water bath for 60 minutes, and 8ml was taken every 5 minutes for MNase-Seq, 10ml for DNA or RNA profiling and 500μl for flow cytometry.

### Flow cytometry of yeast with Sodium Citrate buffer

500μl of yeast culture was spun down, then fixed in 500μl of cold 70% ethanol for 2 hours at room temperature (∼20°C) or overnight at 4°C. After fixation, cells were centrifuged at 13300 rpm for 2 minutes and the pellet was washed with 1ml of 50 mM sodium citrate. Cells were then centrifuged and resuspended in 1ml of 50 mM sodium citrate with 10 μg/ml of RNase and incubated at 37°C for 4 hours. After the RNase treatment, cells were centrifuged and resuspended in 50 mM HCl with 5 mg/ml of pepsin and incubated at 37°C for 30 minutes.

Cells were then washed with 1ml of 50 mM sodium citrate. After centrifugation, cells were resuspended in 1ml of 50 mM Tris pH 7.4 with 0.5 μg/ml of propidium iodide (PI). Finally, tubes were vortexed and 100μl was added to FACS tubes with 1ml of 50 mM Tris pH 7.4 with 0.5 μg/ml of PI. Before processing in the cytometer, cells were sonicated for 8 seconds at 40% amplitude.

### Collection of samples for whole-genome sequencing (replication profiles)

Yeast genomic DNA was extracted using the smash and grab method (https://fangman-brewer.genetics.washington.edu/smash-n-grab.html). DNA was sonicated using the Bioruptor Pico sonicator (Diagenode), and the libraries were prepared according to the TruSeq Nano sample preparation guide from Illumina.

### Collection of samples for whole-transcriptome sequencing (RNA-Seq)

Total RNA was extracted using the RiboPure RNA Purification Kit (Thermo Fisher). Quality and integrity of the RNA was evaluated using an Agilent Tapestation. Libraries were prepared from RNA samples exceeding a RIN quality score of 9.0, according to the TruSeq RNA Library Prep Kit v2 sample preparation guide from Illumina.

### GFP experiments

The *sml1*Δ and *sml1*Δ *SSDDCS thi12Δ::GFP S.cerevisiae* strains were grown as previously described in the block and release experiment, with the modification that cultures were maintained in the 30°C shaking water bath for 150 minutes. Samples were then plated onto 35-mm glass bottom plates (MatTek) precoated with Concanavalin A (Sigma). After 5 min, the medium was changed to SC medium and cells were imaged on a Deltavision widefield fluorescent microscope (GE Healthcare), based on an inverted fluorescence microscope (IX70; Olympus) with an oil immersion Plan-Apochromat 60× NA 1.4 lens (Olympus) for imaging of live cells. Images were acquired, deconvoluted, and projected using SoftWoRx (GE Healthcare). Cells with GFP signal were measured using FIJI.

### Digestion of chromatin with MNase for mono-nucleosome analysis (adapted from [37])

**Day 1** - 8ml of yeast culture collected in each time-point was centrifuged for 2 minutes at 4000 rpm and resuspended in 40ml of 1x PBS, 1% formaldehyde. Samples were mixed and left shaking gently for 10 minutes at room temperature on gyro-rocker to crosslink DNA and proteins. Crosslinking was quenched by adding 5ml of 2.5M glycine and left shaking for 10 mins on orbital shaker at room temperature. Samples were left on ice until all samples have been collected. Then, samples were spun for 5 min at 3200 rpm in 50 ml tubes and washed with 50 ml of sterile ddH2O. Pellets were resuspended and transferred to 1.7ml Axygen tubes, and spun down at top speed in table top centrifuge. Liquid was carefully aspirated and pellets vortexed. Pellets were resuspended in 950μL of zymolyase digestion buffer (ZDB: 50 mM Tris Cl at pH 7.5, 1 M sorbitol, 10 mM β-mercaptoethanol) to remove the cell wall. Then, 100 μL of freshly prepared zymolyase solution (10mg/ml dissolved in ZDB) was added to each sample, and digestion was performed for 60 min at 30°C shaking gently in water bath. Efficiency of digestion was assessed by checking cell morphology under the microscope: cells with digested cell walls will appear spherical. A second test is to take 1μl and dilute to 20μl with ddH20. As cells no longer have a cell wall the osmotic shock will burst them. So absence of cells means zymolyase treatment was successful (spheroplasting). Spheroplasts were pelleted in a microfuge, at 5000 rpm for 5 minutes at 4°C and washed with 1 ml of ZDB. Pellets were then resuspended in 1 ml of spheroplast digestion buffer (SDB: 1 M sorbitol, 50 mM NaCl, 10 mM Tris at pH 8, 5 mM MgCl2, 1 mM CaCl2, 1 mM b-mercaptoethanol, 0.15% NP40). Samples were spun down in the microcentrifuge and gently resuspended in 0.5 ml of SDB. Then 90U of MNase (9μl of 10U/μl MNase solution) was added, and tubes were well mixed and left with gentle agitation for 3 min at 37°C. The amount of MNase needs to be experimentally determined by titration with every batch of MNase. MNase digestion was stopped with the addition of 50 μl of 0.5 M EGTA. Tubes were vortexed after adding EGTA. Samples were treated with RNAse by adding 2 μl of RNase I (100units/μL) for at least 1 hour at 37°C. 10 μl of freshly made up stock of 10mg/ml Proteinase K was added and samples were left at 42°C for at least 3 hours. The formaldehyde cross-links were reversed by incubating samples for > 6 h at 65°C.

**Day 2** - samples were transferred to 2ml rubber-sealed screw-cap tubes. 1 volume of Phenol-Chloroform pH 8 (∼570μl) was added to samples and vortexed and spun for 5 minutes. Aqueous phases were collected to new 2ml lo-bind Eppendorf tube, 5 μl of glycogen and 190 μL of 3M sodium acetate were added and samples were ethanol-precipitated with 1250 μL of cold absolute ethanol (2.5x) and vortexed. Samples were incubated at -20°C overnight and centrifuged at 13000 rpm for 30 minutes at 4°C.

**Day 3 -** Pellets were gently washed by adding 1ml of freshly made 70% ethanol. Then pellets were pulsed down quickly and most volume was carefully aspirated, the wash was repeated with 1ml of 70% ethanol and then tubes were spun down for 15 min at 4°C. Ethanol was carefully aspirated and pellets were air dried for 15 minutes at room temperature, 100μl of Illumina Resuspension Buffer was added and samples incubated for 1 hour at 37°C to redissolve DNA. Size range and relative molarity were determined on a Tapestation using D1000 and Genomic screen tape and total yield was quantified using Qubit broad range dsDNA kit.

### Digestion of chromatin with MNase for transcription factor binding analysis

Same protocol as the one used for mono-nucleosome analysis described above, with the following modifications in the MNase digestion step: after spheroplasting and resuspension in SDB, 5U of MNase (5μl of 1U/μl MNase solution) was added and tubes were incubated on the bench (room temperature) for 20 minutes.

### Library preparation – MNase-Seq mono-nucleosome analysis

250ng of MNase-digested DNA from each sample was end-repaired using the Illumina TruSeq DNA nano kit. AMPure XP beads were added (1.8x volume of DNA, DNA >100bp on beads, <100bp in supernatant) to each reaction to purify the mononucleosomal fragments. A-tailing and adapter ligation was performed using the Illumina TruSeq DNA nano kit. Two subsequent steps of beads purification (1.4x volume of DNA) were performed in order to remove adapter dimers. Based on tests using hyperladder V (25bp bands) the beads can selectively retain DNA of ∼270bp (mono-nucleosomal + adapters) from free adapters (60-120bp) if used at a 1.4x ratio to the volume of DNA sample. PCR cycle quantitation was performed for each sample using KAPA Syber Fast reagents and libraries were PCR amplified using the Illumina TruSeq DNA nano kit, followed by another step of bead purification (1.4x volume of DNA). Library quality and quantity were validated on a Tapestation using D1000 screen tape, Qubit broad range dsDNA kit and NEBNext library quantitation kit for Illumina. Libraries were pooled to final 100nM molarity and one step of bead purification (1x volume of DNA) was performed to completely remove adapter dimers. Finally, 20μl of the pooled libraries at a final 20nM molarity was sequenced in a Illumina HiSeq 1500 platform by the Gurdon Institute Core NGS sequencing facility using 50 bp paired-end reads.

### Library preparation – subMNase-Seq transcription-factor reads

Same as previous with the following modifications: after end-repair and before A-tailing, samples were cleaned by performing a phenol-chloroform precipitation, in order to reduce the volume of the samples without using AMPure XP beads. For adapter ligation, 20% of the amount recommended by Illumina was used (adapters were diluted 1:10 in RSB), in order to minimize adapter dimer formation. This was done because adapter dimers cannot be removed using AMPure XP beads as their size is very similar to TF binding fragments, and smaller fragments are preferentially amplified during library preparation, which means we would be wasting sequencing depth with adapters. After ligation, samples were cleaned using 1.8x AMPure XP beads (DNA >100bp on beads), so TF binding events corresponding to 10-80bp footprint + two adapters (120bp) will bind to the beads.

### Quality control and Mapping

Sample quality was assessed using FastQC High Throughput Sequence QC Report version 0.11.4. All samples were mapped using bowtie2 (version 2.2.6) to the budding yeast reference genome (strain S288C, version R64-2-1), which was indexed using bowtie2-build. SAM files were then converted to BAM, sorted and indexed using samtools (version 0.1.19). The quality control of the alignments was assessed using Qualimap (version 2.2.1).

### Replication profiles

Before generating the replication profiles, sequencing depth was normalised for each timepoint using a bulk value derived from the fraction of the genome that has been replicated at that timepoint (a value between 1 and 2). These values were derived using fitSigmoid https://dzmitry.shinyapps.io/flowfit/.

To generate replication timing profiles, the ratio of uniquely mapped reads in the replicating samples to the non-replicating samples was calculated following [28]. Then, this ratio was plotted for each time-point, and a sigmoid line was fitted. T_rep_ was determined as the time of half-maximal replication (ratio = 1.5). Replication profiles were generated by plotting Trep values for each chromosome location using ggplot2 and smoothed using a moving average in R. All downstream analysis was performed in R.

### RNA-Seq analysis

Read counts for each gene were extracted using genomic ranges and differential expression analysis was performed using DESeq2 [32]. Pair-wise analysis of each time-point was performed using the Wald test. For the time-course analysis, the likelihood ratio test (LRT) was used. Genes were considered differentially expressed if the p-value adjusted value from these tests was < 0.01. PCA analysis was performed using the variance stabilisation transformation (vst command from DESeq2) and plotted using ggplot2. For the k-means clustering, the gene expression data was normalised by row using the scale command and 6 clusters were generated using the k-means command with a maximum of 50 iterations. Heatmaps were generated in R using heatmap.2. Distance to origins, centromeres and telomeres was calculated using HOMER (v4.10.1) [57]. Statistical analyses were performed using R.

### Mono-nucleosome MNase-Seq analysis

Nucleosome calls were identified by processing the BAM files using DANPOS (version 2.2.2) [58]: danpos dpos was used to generate wig files. danpos profile was used to generate the files required for the nucleosome profiles, which were plotted in R. The files with the genomic coordinates of the locations where the heatmaps should be centred were generated using USCS Genome Browser http://genome.ucsc.edu/cgi-bin/hgTables. Peaks in these profiles represent nucleosome dyads and valleys linker DNA or nucleosome depleted regions.

The +1 nucleosome was identified by calculating the distance of nucleosomes to promoters using HOMER. Nucleosomes within -20bp to 80bp of the TSS were classified as +1. The +1 relative position to the TSS was calculated by subtracting the genomic position of the nucleosome to the TSS of the corresponding gene. ACF analysis was performed following [39]. The autocorrelation function was used to determine the pattern of organisation of the first four nucleosomes (+1, +2, +3 and +4) within each gene, using the nucleosome sized reads between 140bp and 180bp overlapping each gene.

### Identification of TF binding motifs genome-wide

To identify TF binding regions genome-wide, the sequence motif was extracted in meme format from the JASPAR database [42], which was used as an input to FIMO [43] to identify all genomic locations where this motif is present. These regions were annotated to genes using HOMER.

### Sub-nucleosomal MNase-Seq analysis

BAM files were converted into bigWig files for visualisation of the results using IGV. For this purpose, bamCoverage (version 3.0.2) was used with the following argument: --binSize 1 -- minFragmentLenght 0 –maxFragmentLength 100.

Identification of high confidence sub-nucleosomal peaks and calculation of fold-change differences between the strains was performed following Gutiérrez et al.: DNA fragments with less than 100bp (sub-nucleosomal events) were selected and the same number of reads was sampled for each fragment size using the time-point with the minimum number of reads for that fragment size. Then, all samples were merged to call all possible peaks in the data. A peak was considered high confidence if the sum of reads mapping to that peak (log2 normalised) across all samples was higher than 75. The log2 ratio of normalised reads occupying each peak between the two strains was calculated for each time-point. Sub-nucleosomal peaks were annotated to TSS and TF binding sites using HOMER as described previously.

### Availability of data and materials

All sequencing data has been deposited in GEO under the accession code GSE199450. All materials are available on request.

## Supplementary Figure Legends

**Supplementary Figure 1.**
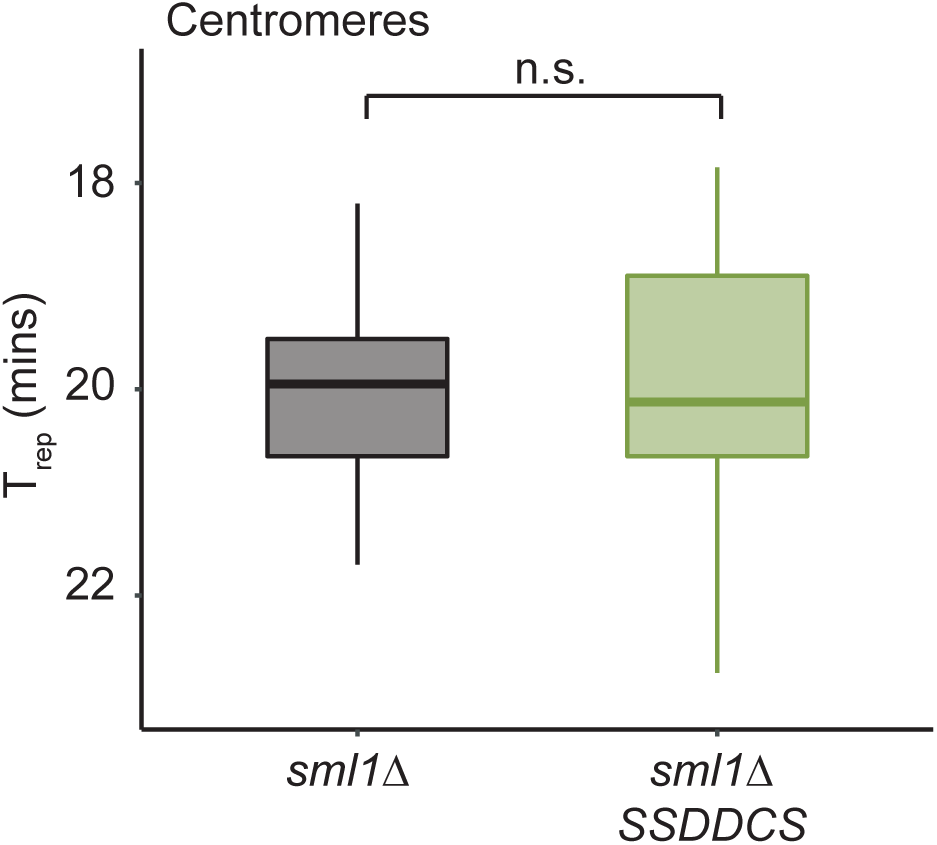
Centromere T_rep_ is unaffected in the SSDDCS strain. Box and whisker plot of the T_rep_ values of the 16 yeast centromeres. Centromeres remained early replicated and were not significantly affected by SSDDCS over-expression (n.s. – non significant, p-value = 0.9578, Welch Two Sample t-test).

**Supplementary Figure 2.**
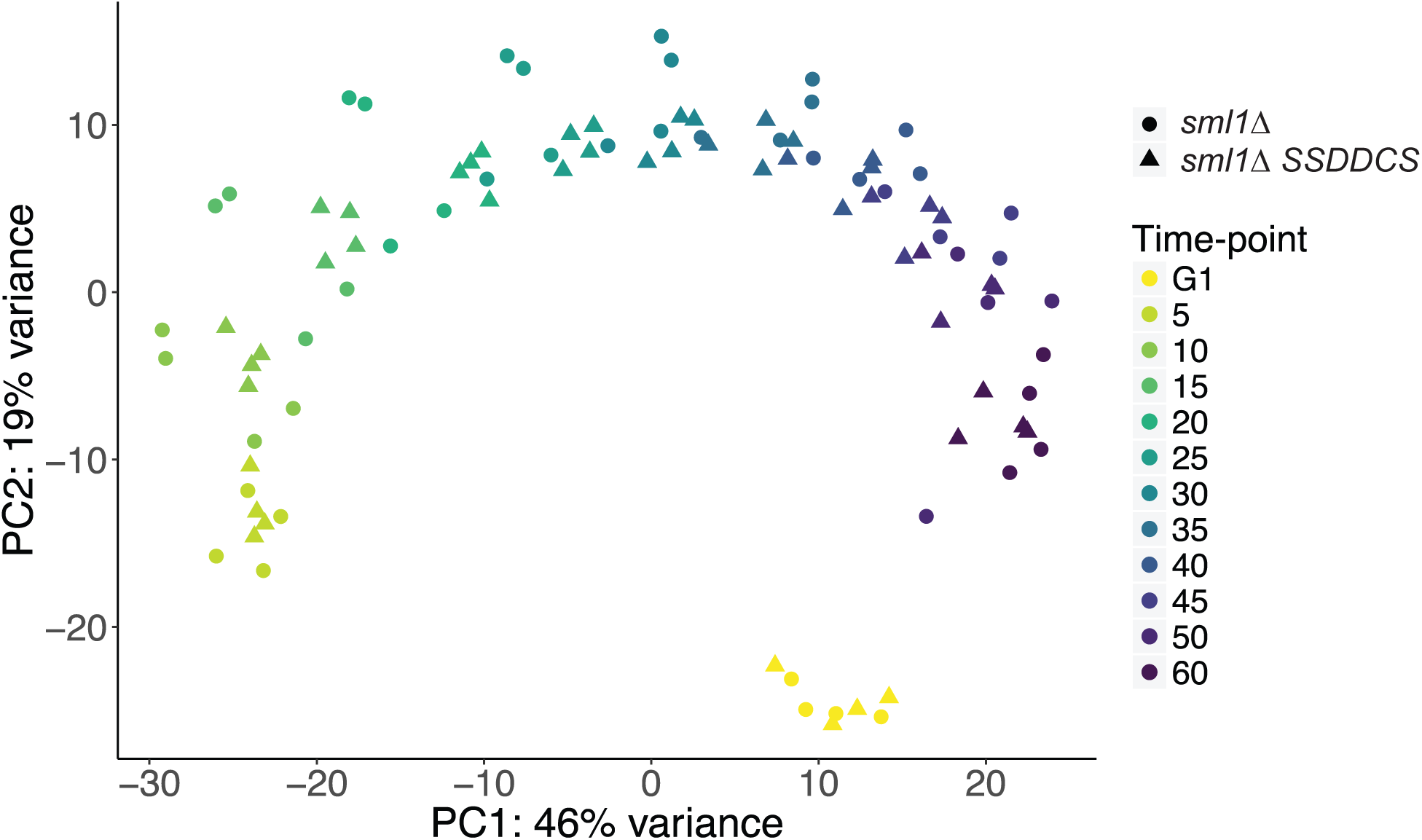
PCA analysis of batch effects in the RNA-seq datasets. Principal component analysis (PCA) plot illustrating the clustering of individual replicates per time-point and strain. Samples are mostly clustered by time-point, illustrating the cell cycle nature of gene expression regulation in budding yeast n = 4.

**Supplementary Figure 3.**
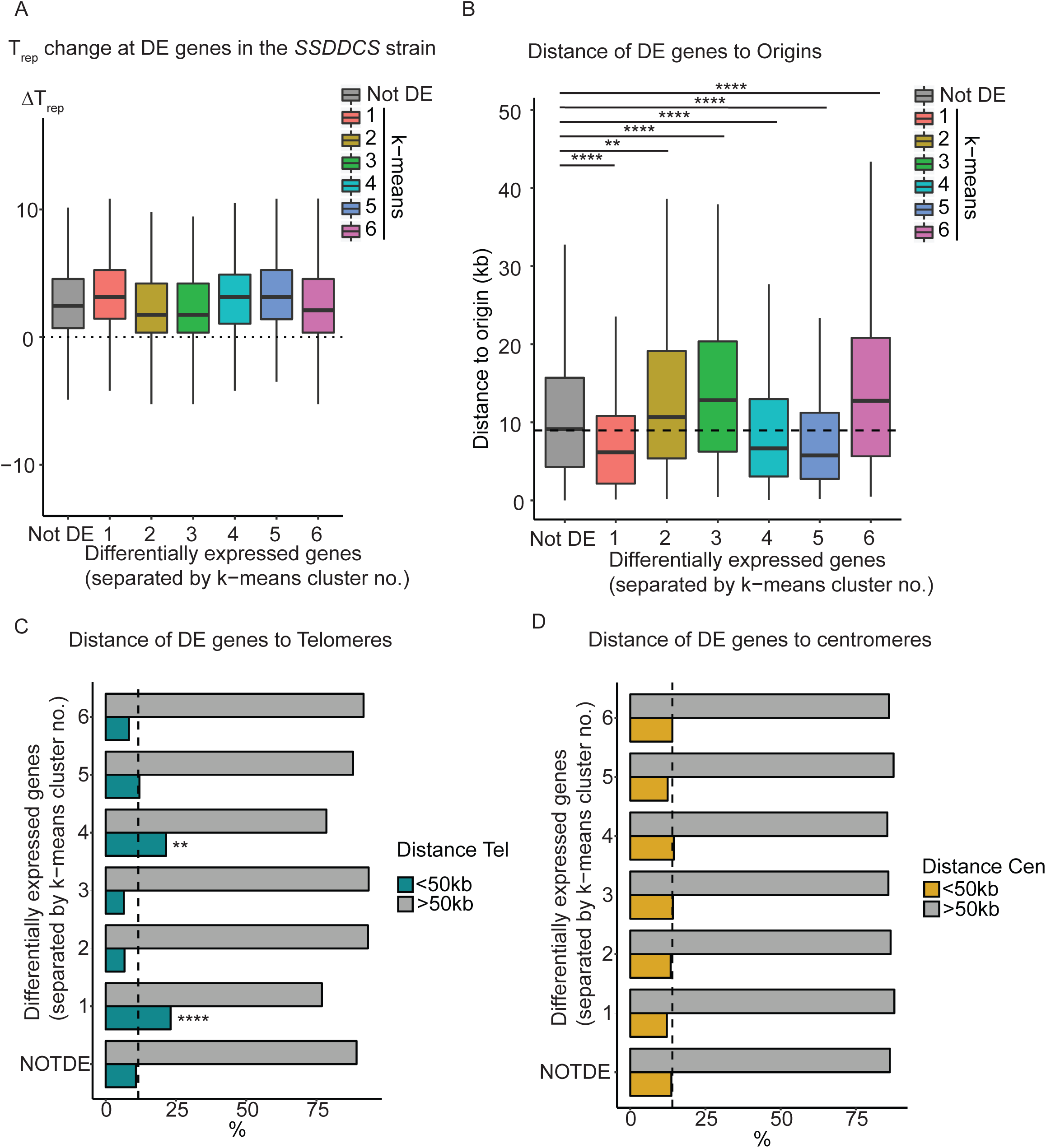
Analysis of the proximity of differentially expressed genes (DE) to origins, telomeres and centromeres. A) Distribution of ΔT_rep_ (*sml1*Δ - *sml1Δ SSDDCS*) for all genes, divided into the k-means clusters from Figure 2C. The dotted line marks 0 (no difference in T_rep_). B) Distribution of gene distances to the closest origin for every gene in the genome, split by k-means cluster. NOT DE represents all non-differentially expressed genes. Dashed horizontal line marks the genome-wide median gene distance to the closest origin = 8.9 kb. p-values are from pairwise comparisons of each k-means cluster versus the non-DE genes using Wilcoxon rank sum test. **** p < 0.0001, ** p < 0.01. C) Proportion of genes which are within or without sub-telomeric regions (less or more than 50kb away from the closest telomere, respectively). Vertical dashed line marks the percentage of all genes located in sub-telomeric regions = 12%. p-values are from an exact binomial test comparison to non-DE genes. **** p < 0.0001, ** p < 0.01. D) Proportion of genes which are within or without sub-centromeric regions (less or more than 50kb away from the centromere, respectively). Vertical dashed line marks the percentage of all genes located in sub-centromeric regions = 14%. All groups had the expected proportion of genes in sub-centromeric regions.

**Supplementary Figure 4.**
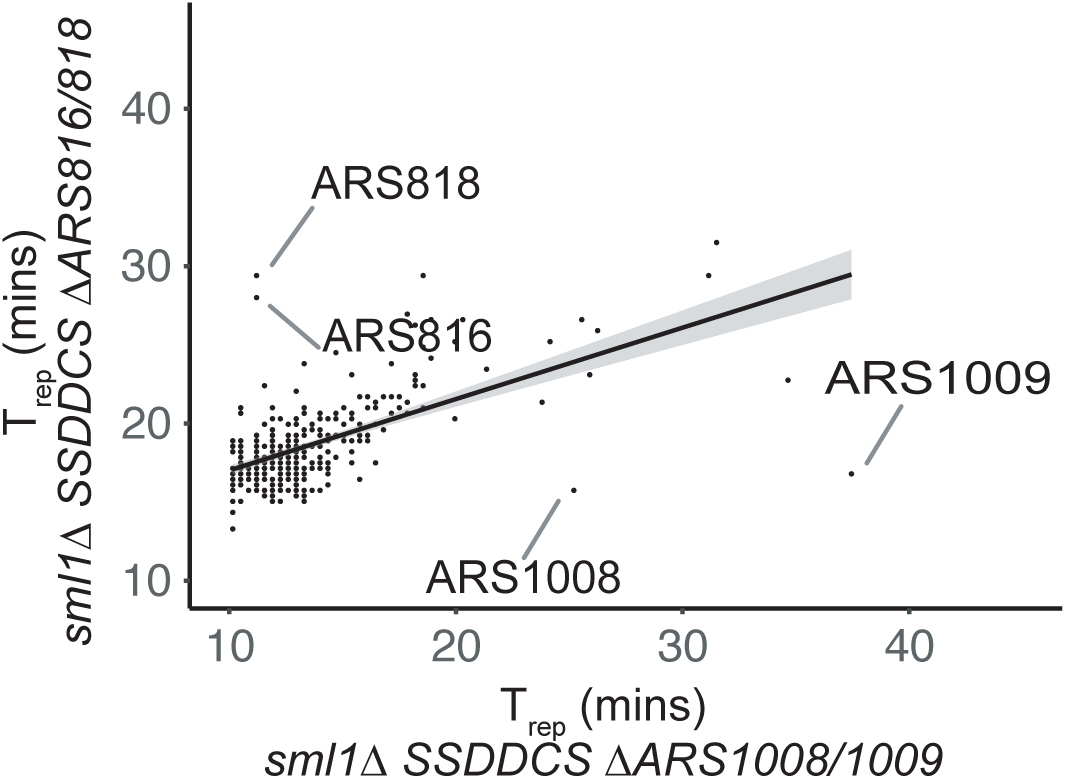
Analysis of the T_rep_ for all origins between the origin deletion strains. Comparison of the T_rep_ for all origins between the *sml1Δ SSDDCS ΔARS816/818* strain (y-axis) and the *sml1Δ SSDDCS ΔARS1008/1009* strain (x-axis). This graph shows that the mutated origins deviate the most from the best fit line for all origins, indicating that the other origins have a similar T_rep_ when comparing between these strains.

## References

1. Bell SP, Labib K: Chromosome Duplication in Saccharomyces cerevisiae. Genetics 2016, 203:1027–1067.

2. Rivera-Mulia JC, Gilbert DM: **Replicating Large Genomes: Divide and Conquer**. Mol Cell 2016, 62:756–765.

3. Muller CA, Nieduszynski CA: Conservation of replication timing reveals global and local regulation of replication origin activity. Genome Res 2012, 22:1953–1962.

4. Ryba T, Hiratani I, Lu J, Itoh M, Kulik M, Zhang J, Schulz TC, Robins AJ, Dalton S, Gilbert DM: Evolutionarily conserved replication timing profiles predict long-range chromatin interactions and distinguish closely related cell types. Genome Res 2010, 20:761–770.

5. Kermi C, Lo Furno E, Maiorano D: Regulation of DNA Replication in Early Embryonic Cleavages. Genes (Basel) 2017, 8.

6. Rivera-Mulia JC, Buckley Q, Sasaki T, Zimmerman J, Didier RA, Nazor K, Loring JF, Lian Z, Weissman S, Robins AJ, et al: **Dynamic changes in replication timing and gene expression during lineage specification of human pluripotent stem cells**. Genome Res 2015, 25:1091–1103.

7. Rivera-Mulia JC, Sasaki T, Trevilla-Garcia C, Nakamichi N, Knapp D, Hammond CA, Chang BH, Tyner JW, Devidas M, Zimmerman J, et al: **Replication timing alterations in leukemia affect clinically relevant chromosome domains**. Blood Adv 2019, 3:3201–3213.

8. Raghuraman MK, Brewer BJ, Fangman WL: Cell cycle-dependent establishment of a late replication program. Science 1997, 276:806–809.

9. Dimitrova DS, Gilbert DM: The spatial position and replication timing of chromosomal domains are both established in early G1 phase. Mol Cell 1999, 4:983–993.

10. Rhind N, Gilbert DM: **DNA replication timing**. Cold Spring Harb Perspect Biol 2013, 5:a010132.

11. Ferguson BM, Fangman WL: A position effect on the time of replication origin activation in yeast. Cell 1992, 68:333–339.

12. Yoshida K, Bacal J, Desmarais D, Padioleau I, Tsaponina O, Chabes A, Pantesco V, Dubois E, Parrinello H, Skrzypczak M, et al: The histone deacetylases sir2 and rpd3 act on ribosomal DNA to control the replication program in budding yeast. Mol Cell 2014, 54:691–697.

13. Vogelauer M, Rubbi L, Lucas I, Brewer BJ, Grunstein M: **Histone acetylation regulates the time of replication origin firing**. Mol Cell 2002, 10:1223–1233.

14. Belsky JA, MacAlpine HK, Lubelsky Y, Hartemink AJ, MacAlpine DM: Genome-wide chromatin footprinting reveals changes in replication origin architecture induced by pre-RC assembly. Genes Dev 2015, 29:212–224.

15. Hoggard T, Shor E, Muller CA, Nieduszynski CA, Fox CA: A Link between ORC-origin binding mechanisms and origin activation time revealed in budding yeast. PLoS Genet 2013, 9:e1003798.

16. Dukaj L, Rhind N: The capacity of origins to load MCM establishes replication timing patterns. PLoS Genet 2021, 17:e1009467.

17. Tanaka S, Nakato R, Katou Y, Shirahige K, Araki H: Origin association of Sld3, Sld7, and Cdc45 proteins is a key step for determination of origin-firing timing. Curr Biol 2011, 21:2055–2063.

18. Mantiero D, Mackenzie A, Donaldson A, Zegerman P: Limiting replication initiation factors execute the temporal programme of origin firing in budding yeast. EMBO J 2011, 30:4805–4814.

19. Collart C, Allen GE, Bradshaw CR, Smith JC, Zegerman P: Titration of four replication factors is essential for the Xenopus laevis midblastula transition. Science 2013, 341:893–896.

20. Fang D, Lengronne A, Shi D, Forey R, Skrzypczak M, Ginalski K, Yan C, Wang X, Cao Q, Pasero P, Lou H: Dbf4 recruitment by forkhead transcription factors defines an upstream rate-limiting step in determining origin firing timing. Genes Dev 2017, 31:2405–2415.

21. Natsume T, Muller CA, Katou Y, Retkute R, Gierlinski M, Araki H, Blow JJ, Shirahige K, Nieduszynski CA, Tanaka TU: **Kinetochores coordinate pericentromeric cohesion and early DNA replication by Cdc7-Dbf4 kinase recruitment**. Mol Cell 2013, 50:661–674.

22. Richards L, Das S, Nordman JT: **Rif1-Dependent Control of Replication Timing**. Genes (Basel) 2022, 13.

23. Lima-de-Faria A, Jaworska H: **Late DNA synthesis in heterochromatin**. Nature 1968, 217:138–142.

24. Rivera-Mulia JC, Gilbert DM: Replication timing and transcriptional control: beyond cause and effect-part III. Curr Opin Cell Biol 2016, 40:168–178.

25. . Fraser HB: Cell-cycle regulated transcription associates with DNA replication timing in yeast and human. Genome Biol 2013, 14:R111.

26. Klein KN, Zhao PA, Lyu X, Sasaki T, Bartlett DA, Singh AM, Tasan I, Zhang M, Watts LP, Hiraga SI, et al: **Replication timing maintains the global epigenetic state in human cells**. Science 2021, 372:371–378.

27. Muller CA, Nieduszynski CA: DNA replication timing influences gene expression level. J Cell Biol 2017, 216:1907–1914.

28. Batrakou DG, Muller CA, Wilson RHC, Nieduszynski CA: DNA copy-number measurement of genome replication dynamics by high-throughput sequencing: the sort-seq, sync-seq and MFA-seq family. Nat Protoc 2020, 15:1255–1284.

29. McGuffee SR, Smith DJ, Whitehouse I: Quantitative, genome-wide analysis of eukaryotic replication initiation and termination. Mol Cell 2013, 50:123–135.

30. Siow CC, Nieduszynska SR, Muller CA, Nieduszynski CA: **OriDB, the DNA replication origin database updated and extended**. Nucleic Acids Res 2012, 40:D682–686.

31. Morafraile EC, Hanni C, Allen G, Zeisner T, Clarke C, Johnson MC, Santos MM, Carroll L, Minchell NE, Baxter J, et al: **Checkpoint inhibition of origin firing prevents DNA topological stress**. Genes Dev 2019, 33:1539–1554.

32. Love MI, Huber W, Anders S: Moderated estimation of fold change and dispersion for RNA-seq data with DESeq2. Genome Biol 2014, 15:550.

33. Spellman PT, Sherlock G, Zhang MQ, Iyer VR, Anders K, Eisen MB, Brown PO, Botstein D, Futcher B: Comprehensive identification of cell cycle-regulated genes of the yeast Saccharomyces cerevisiae by microarray hybridization. Mol Biol Cell 1998, 9:3273–3297.

34. Lu Z, Lin Z: Pervasive and dynamic transcription initiation in Saccharomyces cerevisiae. Genome Res 2019, 29:1198–1210.

35. Voichek Y, Bar-Ziv R, Barkai N: **Expression homeostasis during DNA replication**. Science 2016, 351:1087–1090.

36. Osborne EA, Hiraoka Y, Rine J: Symmetry, asymmetry, and kinetics of silencing establishment in Saccharomyces cerevisiae revealed by single-cell optical assays. Proc Natl Acad Sci U S A 2011, 108:1209–1216.

37. Nocetti N, Whitehouse I: Nucleosome repositioning underlies dynamic gene expression. Genes Dev 2016, 30:660–672.

38. Jiang C, Pugh BF: A compiled and systematic reference map of nucleosome positions across the Saccharomyces cerevisiae genome. Genome Biol 2009, 10:R109.

39. Gutierrez MP, MacAlpine HK, MacAlpine DM: Nascent chromatin occupancy profiling reveals locus- and factor-specific chromatin maturation dynamics behind the DNA replication fork. Genome Res 2019, 29:1123–1133.

40. Lenstra TL, Benschop JJ, Kim T, Schulze JM, Brabers NA, Margaritis T, van de Pasch LA, van Heesch SA, Brok MO, Groot Koerkamp MJ, et al: **The specificity and topology of chromatin interaction pathways in yeast**. Mol Cell 2011, 42:536–549.

41. Ramachandran S, Henikoff S: Transcriptional Regulators Compete with Nucleosomes Post-replication. Cell 2016, 165:580–592.

42. Sandelin A, Alkema W, Engstrom P, Wasserman WW, Lenhard B: **JASPAR: an open-access database for eukaryotic transcription factor binding profiles**. Nucleic Acids Res 2004, 32:D91–94.

43. Grant CE, Bailey TL, Noble WS: **FIMO: scanning for occurrences of a given motif**. Bioinformatics 2011, 27:1017–1018.

44. MacIsaac KD, Wang T, Gordon DB, Gifford DK, Stormo GD, Fraenkel E: **An improved map of conserved regulatory sites for Saccharomyces cerevisiae**. BMC Bioinformatics 2006, 7:113.

45. Pak J, Segall J: Regulation of the premiddle and middle phases of expression of the NDT80 gene during sporulation of Saccharomyces cerevisiae. Mol Cell Biol 2002, 22:6417–6429.

46. Fennessy RT, Owen-Hughes T: Establishment of a promoter-based chromatin architecture on recently replicated DNA can accommodate variable inter-nucleosome spacing. Nucleic Acids Res 2016, 44:7189–7203.

47. Stewart-Morgan KR, Petryk N, Groth A: **Chromatin replication and epigenetic cell memory**. Nat Cell Biol 2020, 22:361–371.

48. Omberg L, Meyerson JR, Kobayashi K, Drury LS, Diffley JF, Alter O: Global effects of DNA replication and DNA replication origin activity on eukaryotic gene expression. Mol Syst Biol 2009, 5:312.

49. Hwang Y, Hidalgo D, Socolovsky M: The shifting shape and functional specializations of the cell cycle during lineage development. WIREs Mech Dis 2021, 13:e1504.

50. Rivera-Mulia JC, Schwerer H, Besnard E, Desprat R, Trevilla-Garcia C, Sima J, Bensadoun P, Zouaoui A, Gilbert DM, Lemaitre JM: **Cellular senescence induces replication stress with almost no affect on DNA replication timing**. Cell Cycle 2018, 17:1667–1681.

51. Sati S, Bonev B, Szabo Q, Jost D, Bensadoun P, Serra F, Loubiere V, Papadopoulos GL, Rivera-Mulia JC, Fritsch L, et al: **4D Genome Rewiring during Oncogene-Induced and Replicative Senescence**. Mol Cell 2020, 78:522–538 e529.

52. Hu Z, Chen K, Xia Z, Chavez M, Pal S, Seol JH, Chen CC, Li W, Tyler JK: Nucleosome loss leads to global transcriptional up-regulation and genomic instability during yeast aging. Genes Dev 2014, 28:396–408.

53. Chandra T, Ewels PA, Schoenfelder S, Furlan-Magaril M, Wingett SW, Kirschner K, Thuret JY, Andrews S, Fraser P, Reik W: **Global reorganization of the nuclear landscape in senescent cells**. Cell Rep 2015, 10:471–483.

54. Vastenhouw NL, Cao WX, Lipshitz HD: **The maternal-to-zygotic transition revisited**. Development 2019, 146.

55. Hug CB, Vaquerizas JM: The Birth of the 3D Genome during Early Embryonic Development. Trends Genet 2018, 34:903–914.

56. Seller CA, O’Farrell PH: Rif1 prolongs the embryonic S phase at the Drosophila mid-blastula transition. PLoS Biol 2018, 16:e2005687.

57. Heinz S, Benner C, Spann N, Bertolino E, Lin YC, Laslo P, Cheng JX, Murre C, Singh H, Glass CK: Simple combinations of lineage-determining transcription factors prime cis-regulatory elements required for macrophage and B cell identities. Mol Cell 2010, 38:576–589.

58. Chen K, Xi Y, Pan X, Li Z, Kaestner K, Tyler J, Dent S, He X, Li W: **DANPOS: dynamic analysis of nucleosome position and occupancy by sequencing**. Genome Res 2013, 23:341–351.

